# Dysregulated Skeletal Muscle Myosin Super-relaxation and Energetics in Type II Diabetes

**DOI:** 10.1101/2024.06.12.598615

**Authors:** Christopher T. A. Lewis, Roger Moreno-Justicia, Lola Savoure, Enrique Calvo, Agata Bak, Jenni Laitila, Robert A.E. Seaborne, Steen Larsen, Hiroyuki Iwamoto, Marina Cefis, Jose A. Morais, Gilles Gouspillou, Jorge Alegre-Cebollada, Thomas J. Hawke, Jesús Vazquez, Miquel Adrover, Vincent Marcangeli, Atul S. Deshmukh, Julien Ochala

**Affiliations:** Department of Biomedical Sciences, University of Copenhagen, Copenhagen, Denmark; Novo Nordisk Foundation Center for Basic Metabolic Research, Faculty of Health and Medical Sciences, University of Copenhagen, Denmark; Centro Nacional de Investigaciones Cardiovasculares (CNIC), Madrid, Spain; Centre for Human and Applied Physiological Sciences, Faculty of Life Sciences & Medicine, King’s College London, London, UK; Clinical Research Centre, Medical University of Bilalystok, Bialystok, Poland; Spring-8, Japan Synchrotron Radiation Research Institute, Hyogo, Japan; Département des Sciences de l’Activité Physique, Faculté des Sciences, UQAM, Montréal, Québec, Canada; Department of Medicine, Research Institute of the McGill University Health Centre, Montréal, Québec, Canada; Department of Pathology and Molecular Medicine, McMaster University, Hamilton, Ontario, Canada; Institut Universitari d’Investigació en Ciències de la Salut (IUNICS), Institut d’Investigació Sanitària Illes Balears (IdISBa), Departament de Química, Universitat de les Illes Balears, Palma de Mallorca, Spain; Département des sciences biologiques, Faculté des Sciences, UQAM, Montréal, Québec, Canada; CIBER de Enfermedades Cardiovasculares (CIBERCV), Madrid, Spain

**Keywords:** skeletal muscle, metabolism, diabetes, myosin

## Abstract

Disrupted energy balance is critical for the onset and development of Type II diabetes. The exact underlying metabolic mechanisms remain incomplete but skeletal muscle is thought to play an important pathogenic role. As the super-relaxed state of its most abundant protein, myosin, regulates cellular energetics, here, we aimed to investigate whether it is altered in patients with type II diabetes. For that, we used vastus lateralis biopsy specimens (obtained from patients with type II diabetes and matched controls) and run a combination of structural and functional assays consisting of loaded Mant-ATP chase experiments, X-ray diffraction and LC-MS/MS proteomics in isolated muscle fibres. Our studies revealed a greater muscle myosin super-relaxation and decreased cellular ATP demand in patients than controls. Subsequent proteomic analyses indicated that these (mal)adaptations likely originated from remodeled sarcomeric proteins and greater myosin glycation levels in patients than controls. Overall, our findings emphasize a complex molecular dysregulation of myosin super-relaxed state and energy consumption in type II diabetes. Ultimately, pharmacological targeting of myosin could benefit skeletal muscle and whole-body metabolic health through the enhancement of ATP consumption.

**Significance Statement:** Myosin super-relaxation, essential for the regulation of skeletal muscle metabolic rate, is disrupted in type II diabetes due to protein hyper-glycation. As a consequence, myosin ATP demand is significantly lowered. Overall, our findings provide a strong rationale for the use of activators of myosin ATPase to enhance basal energy expenditure in type II diabetes.

## Introduction

As a major tissue in the regulation of whole-body glycemic control, skeletal muscle has long been established as critical to the onset of type 2 diabetes mellitus (T2DM) (1). Whilst decades of research have advanced our molecular understanding of how skeletal muscle contributes to the etiology of such disease, few of these studies have investigated molecular changes in energy expenditure and whether the most abundant protein, myosin, is involved.

Skeletal muscle contraction is highly energy-dependent, and ATP is consumed directly by the head region of the myosin molecule, which is itself an ATPase (2). Until recently, energy usage was thought to be mostly linked to the consumption of ATP by active myosin molecules. However, this doctrine has been challenged following the discovery that myosin maintains a significant consumption of ATP, and that the rate of this ATP consumption is dependent on its resting state (3). It is now established that myosin can adopt at least two different biochemical states at rest, known as the disordered-relaxed state and the super-relaxed state (3-5). In the disordered-relaxed state (DRX), the myosin heads are likely to be in a structural ON state. They are not bound to actin and can exist freely in the interfilamentous space of the sarcomere (6, 7). In the super-relaxed state (SRX), the head regions of the myosin molecules have a structural OFF conformation in which they are folded backwards against the thick filament backbone of the sarcomere (6, 8). This folding means that in the SRX state, the ATPase site located on the head region of the myosin molecule is sterically inhibited, preventing ATP binding (7, 9). Therefore, myosin molecules which are in SRX, have an ATP turnover rate which is approximately ten times slower than those in DRX (4, 10, 11).

Muscle myosin molecules exist in a carefully controlled ratio of DRX:SRX (12). Dysregulation of this ratio occurs in inherited conditions of both cardiac and skeletal muscle following mutations to genes encoding proteins of the sarcomere (13-15). It has been estimated that a 20% shift of myosin heads from SRX to DRX in skeletal muscle would increase whole-body energy expenditure by 16% (4). Thus, uncovering changes to the DRX:SRX ratio in metabolic diseases, such as T2DM, is of great interest as it would unravel major molecular disturbances in the energy demand. Hence, in the present study we aimed to explore the hypothesis that, in T2DM, there would be a re-modelling of the proportions of myosin molecules in DRX and SRX, ultimately affecting the ATP consumption of resting skeletal muscle.

The main cellular source of ATP is glycolysis, a process that is not harmless for cells, as it produces highly reactive carbonyl compounds as side products, including methylglyoxal (MG). MG has high intrinsic reactivity making it one of the endogenous compounds with higher ability to randomly modify nucleophilic groups of long-life biomolecules (16). Due to this, evolution has designed enzymatic mechanisms to control the levels of intracellular MG, such as the glyoxalase system (Glo-1 and Glo-2). However, both enzymes become downregulated in T2DM and consequently, the levels of MG in diabetic patients are 2 to 6-fold higher than in non-diabetic individuals (17). Once it is formed, MG rapidly reacts with Arg, Lys and Cys side-chains producing MG-derived advanced glycation end-products (AGEs), such as N^ε^-(carboxyethyl)lysine (CEL), hydroimidazolones (MG-Hs; which include MG-H1, MG-H2 and MG-H3) or 2-ammonio-6-[1-(5-ammonio-6-oxido-6-oxohexyl)-5-methylimidazolium-3-yl] hexanoate (MOLD) (18, 19). The formation of these AGEs changes the chemical nature of the protein residues, potentially impacting the protein structure and thus their function, potentially exacerbating the progression of T2DM (20). Some recent studies have demonstrated that in patients with T2DM, glycation can also occur on myosin, impacting its function (21, 22). Therefore, we further aimed to test the hypothesis that changes in the myosin DRX:SRX ratio in T2DM would intrinsically be caused by aberrant levels of glycation.

## Results

Our investigations consisted of isolating individual skeletal muscle fibres from the *vastus lateralis* of male patients with T2DM and controls, who were matched for age and body-mass index (Table S1).

### Patients with T2DM have higher levels of myosin SRX/OFF state in type I muscle fibres

To test our hypothesis that myosin DRX to SRX ratio could be altered in patients with T2DM, we performed loaded Mant-ATP chase assays on isolated and permeabilized single muscle fibres (Fig. 1a). A total of 284 individual myofibres were tested (between 8 and 12 fibres per individual). A fibre-type breakdown from these samples is shown in Table S1. Interestingly, we found a significantly lower number of myosin heads in DRX (P1) in type I fibres of T2DM patients than in controls (Fig. 1b). This finding was matched by a significantly higher percentage of fibres in SRX (P2) in type I fibres of T2DM patients than in controls (Fig. 1c). Despite these, we did not observe any differences between the groups in the ATP turnover times of the myosin molecule in DRX (T1) (Fig. 1d) or SRX (T2) (Fig. 1e). These results demonstrate that in T2DM, resting myosin state undergoes remodeling causing a decrease in the ATP consumption of type I, oxidative, fibres.

**Figure 1.**
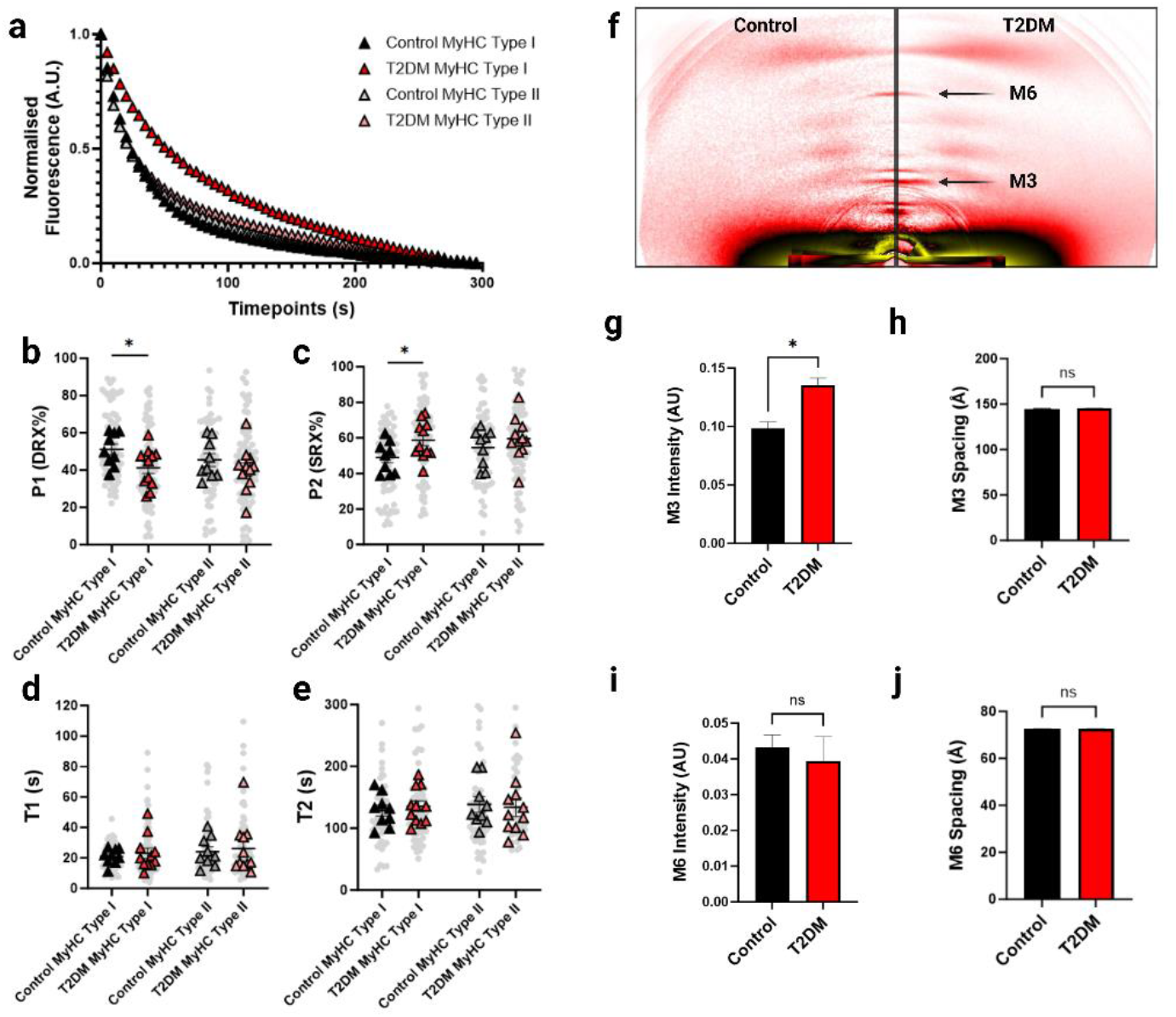
Myosin relaxed states are altered in T2DM. **a**. Representative fluorescence mant-ATP decays from single muscle fibres isolated from control and T2DM skeletal muscle biopsies and measured over 300 seconds. **b-c**. Percentage of myosin heads in the P1/DRX (b) or P2/SRX (c) from single muscle fibres obtained from control and T2DM patients. Values were separated based on each individual fibre was MyHC type I or MyHC type II. **d**. T1 value in seconds denoting the ATP turnover lifetime of the DRX. **e**. T2 value in seconds denoting the ATP turnover lifetime in seconds of the SRX. Grey circles represent the values from each individual muscle fibre which was analyzed. Colored triangles represent the mean value from an individual participant, 8-12 fibres analyzed per participant. Statistical analysis was performed upon mean values. Unpaired student’s t-test was used to calculate statistical significance. n = 9-11 participant per group. **f**. Representative x-ray diffraction recordings from permeabilized skeletal muscle bundles from control and T2DM participant. The M3 and M6 meridional reflections are indicated with black arrows. **g**. Normalized intensity (A.U.) of the M3 meridional reflection. **h**. M3 meridional spacing, measured in ångstrom (Å). **i**. Normalized intensity (A.U.) of the M6 meridional reflection. j. M6 meridional spacing, measured in ångstrom (Å). Data is displayed as mean ± SEM. Differences were tested using a bilateral Students’s t-test. n = 6 participants analyzed per group. * = *p* < 0.05.

To gain further structural insights and relate the above DRX/SRX with myosin ON/OFF states, we performed small-angle X-ray diffraction on the same muscle biopsy specimens and focused on myosin meridional reflections, namely M3 and M6 (Fig. 1f). We found that the M3 intensity was significantly greater in T2DM patients than in controls (Fig. 1g-h). This suggests that, even though the distance between myosin crowns is preserved, there is a higher order of myosin heads along the thick filament (OFF state) (23). We did not observe any change in M6 intensity or spacing between the groups (Fig. 1i-j), highlighting the maintenance of thick filament compliance/extensibility (23). Altogether, our findings support that, in type I fibres of T2DM patients, myosin molecules adopt a preferred ATP-conserving SRX and OFF state without any other major thick filament disturbances.

### The coiled coil domain of the *MYH7* protein is glycated in individuals with T2DM

To get a deeper understanding of the molecular events leading to the changes in myosin resting states we assessed the level of glycation on myosin in these samples. A graphical schematic illustrates the formation of MG and the formation of MG-derived AGEs (Fig. 2a). We identified several AGE-modified peptides arising from the myosin heavy chain type I (*MYH7*) protein obtained from T2DM patients (Fig. S1), which were absent in the controls (Fig. 2b). The identification of these glycated peptides was in-depth validated by a highly sensitive scan using the VseqExplorer application to calculate quantitative E-scores (see Materials and Methods). These glycated peptide fragments were mainly located upon the coiled coil region of the *MYH7* protein (Fig. 2b-c).

**Figure 2.**
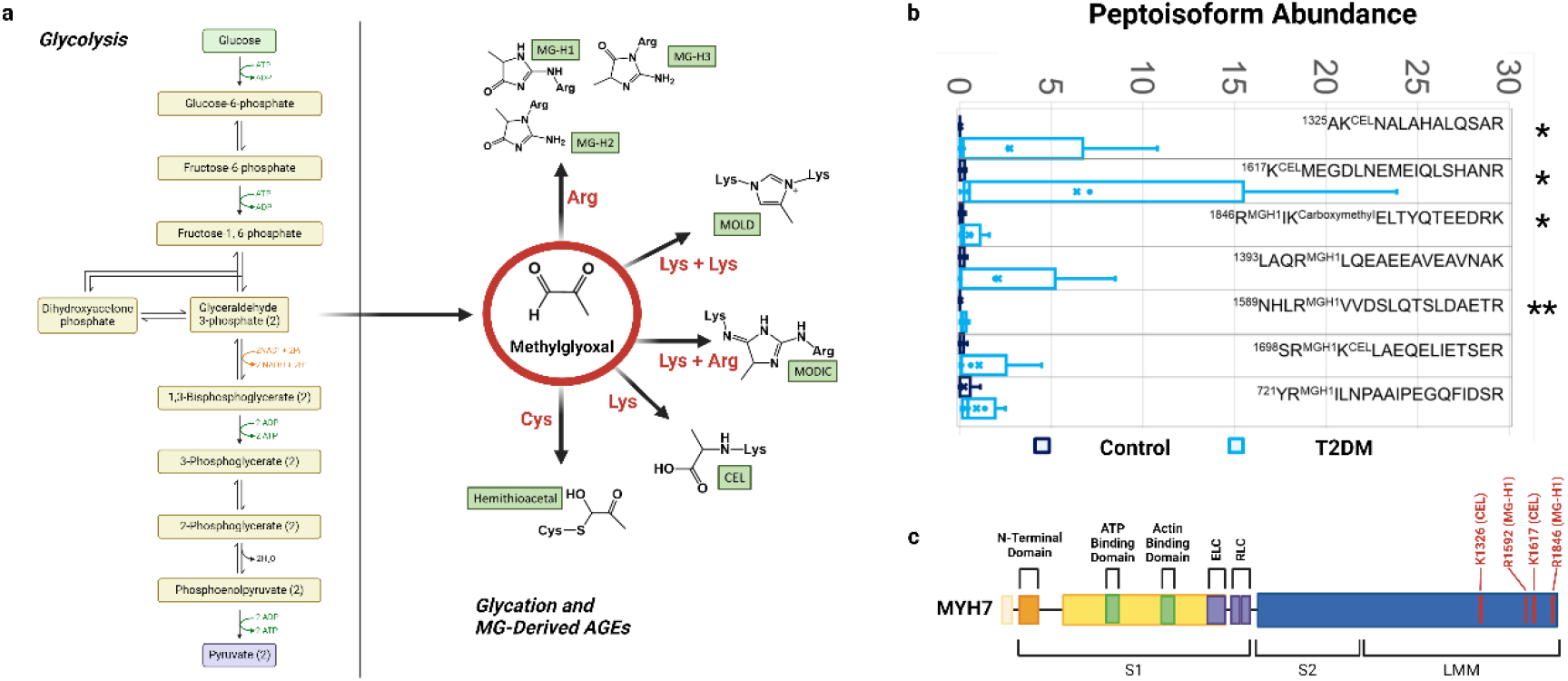
Glycation of type I myosin heavy chain (*MYH7*) in T2DM. **a**. Schematic of the process of glycolysis from which is formed the intracellular MG and the chemical formulae of some of the MG-derived AGEs that have been detected on different protein residues. **b**. Quantification of several glycated peptides from *MYH7* showing increased glycation in T2DM participants when compared with the controls. Mann-Whitney test was used to calculate statistical significance. * = *p* < 0.05, ** = *p* < 0.01. n = 5 individual participants per group. We could detect several peptide fragments modified through the formation of N^ε^-(carboxyethyl)lysine (CEL), hydroimidazolones (MG-Hs) or N^ε^-(carboxymethyl)lysine. **c**. Scheme of *MYH7* protein regions indicating the location of the glycated residues in red. Figure created using BioRender.com.

### Acute glycation of type I muscle fibres induces an increase of the percentage of resting myosin heads in SRX

To emphasize the functional effects of glycation on myosin proteins, we performed paired loaded Mant-ATP chase experiments with and without MG after 30-minute incubation (Fig. 3a). Consistent with our previous results, the baseline percentage of myosin heads in DRX was lower in type I muscle fibres of T2DM patients than in controls (Fig. 3b). This was matched with a greater proportion of SRX in type I muscle fibres of T2DM patients than in controls (Fig. 3c). Following incubation with MG, the fraction of myosin heads in DRX of type I fibres from controls significantly decreased but not of type II fibres (Fig. 3b). Again, this was matched with a significant increase in the amount of SRX in type I fibres of controls but not in type II fibres (Fig. 3c). Conversely, we did not find any change following MG incubation in either fibre type in T2DM patients (Fig. 3b-c). A plausible reason for such discrepancy is likelihood that fibres from T2DM patients were already saturated and hyper-glycated at the time of the baseline experiments. MG incubations also had an impact on the ATP turnover time of the resting myosin molecules (Fig. 3d-e).

**Figure 3.**
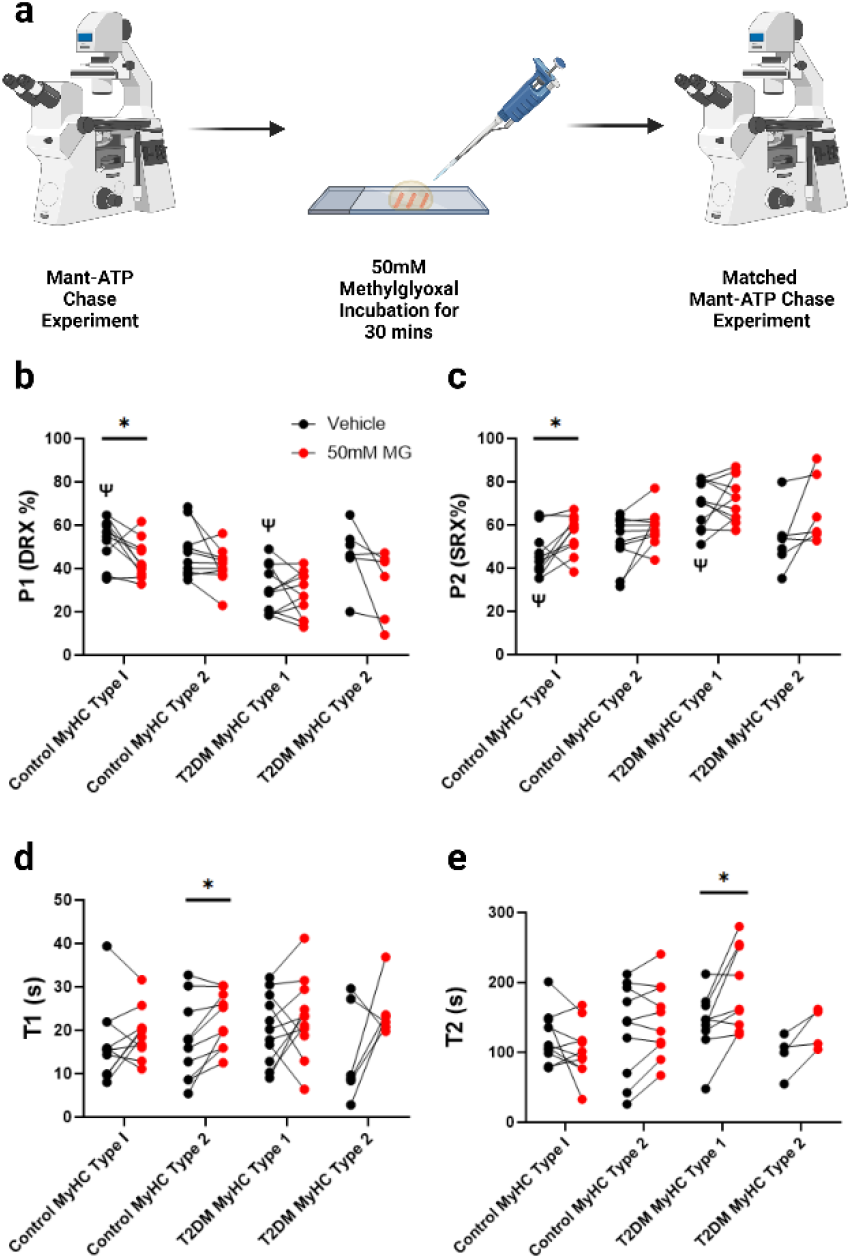
Acute glycation increased the percentage of myosin heads in the SRX in control participants. **a**. Workflow of loaded Mant-ATP chase experiments in which single muscle fibres were incubated with 50mM MG in between matched experimental runs. Figure created using Biorender.com **b**. Percentage of myosin heads in the P1/DRX before (black circles) and after (red circles) treatment with MG. **c**. or Percentage of myosin heads in the P2/SRX before (black circles) and after (red circles) treatment with MG. Values were separated based on each individual fibre was MyHC type I or MyHC type II. **d**. T1 value in seconds denoting the ATP turnover lifetime of the DRX before (black circles) and after (red circles) treatment with MG. **e**. T2 value in seconds denoting the ATP turnover lifetime in seconds of the SRX before (black circles) and after (red circles) treatment with MG. Values were separated based on each individual fibre was MyHC type I or MyHC type II. Paired Student’s t-test was used to calculate statistical significance. * = *p* < 0.05 between fibres before and after MG incubation, Ψ = *p* < 0.05 between groups. n = 5 participants per group.

### Differential expression of sarcomeric proteins in type I muscle fibres of patients with T2DM

Besides post-translational modifications and glycation, other more profound remodelling may occur in T2DM and may include alterations in skeletal muscle protein expression. To investigate this, we performed high-throughput single muscle fibre proteomics using the same muscle biopsy specimens (Fig. 4a). A total of 109 single muscle fibres were analyzed from 10 different muscle biopsies (5 per group). On an average 584 proteins were quantified in each single muscle fibre (Table S2). The samples were grouped by fibre type as determined by immunohistochemistry, which was validated by the proteomics data. The validation revealed expected enrichment of known type I or II fibre specific proteins, such as TNNT1 and TNNI1 or MYH2 and MYBPC2 respectively (Fig. 4b). This validation of the method via the examination of expression profiles between fibre types provided a high level of confidence to proceed and look at the specific changes between the control and T2DM groups in each individual fibre type.

**Figure 4.**
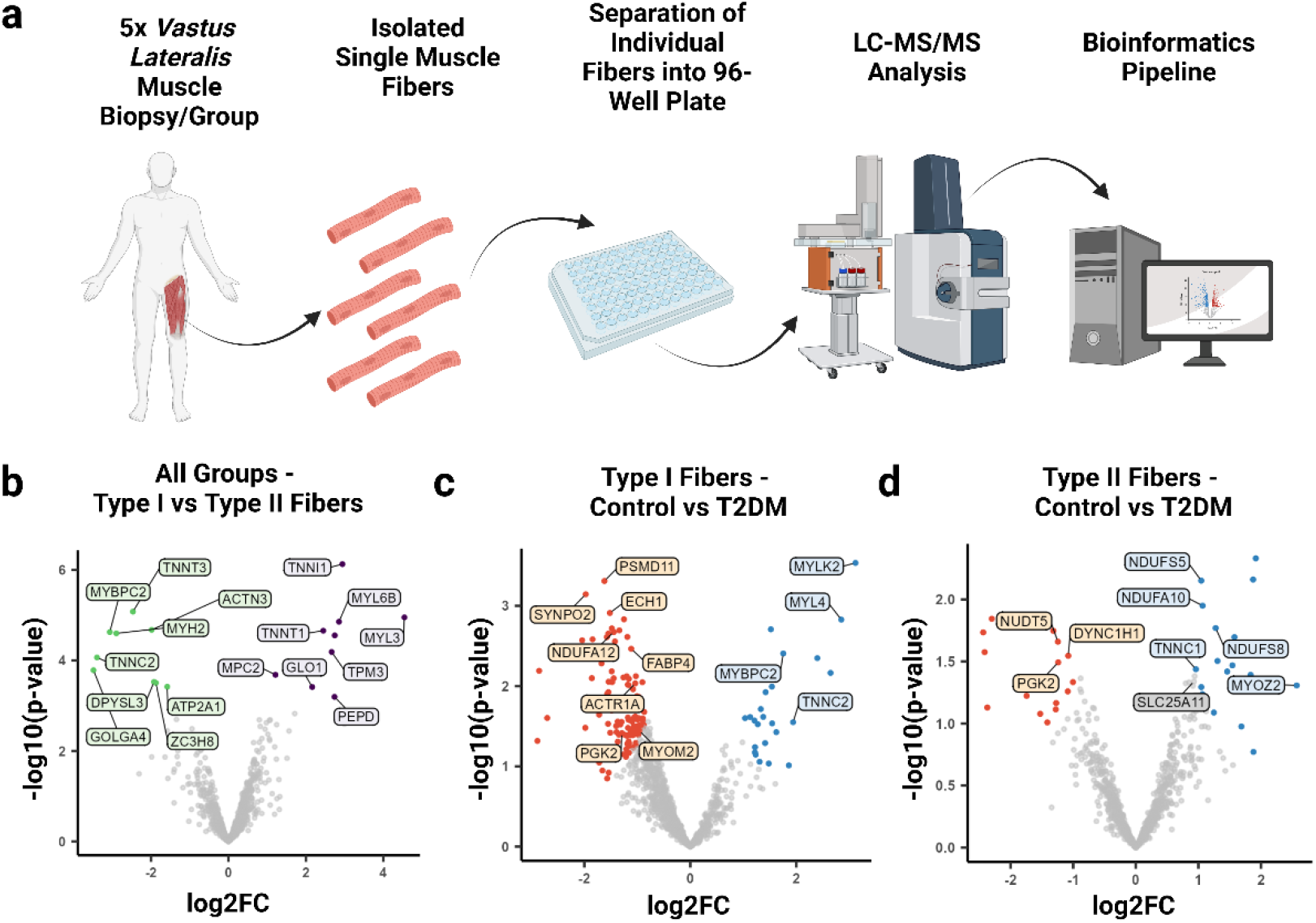
Single fibre proteomics shows that type I muscle fibres from T2DM patients have differential expression of sarcomeric proteins. **a**. Workflow of the isolation of single skeletal muscle fibres from *vastus lateralis* muscle biopsies and downstream processing of these single muscle fibres using mass spectrometry and subsequent bioinformatics pipeline. Figure created using Biorender.com. **b**. Volcano plot demonstrating differentially expressed proteins between type I and type II muscle fibres from both participant groups. **c**. Volcano plot showing proteins that are differentially expressed in type I fibres between control and T2DM groups. **d**. Volcano plot showing proteins that are differentially expressed in type II fibres between control and T2DM groups. Detailed information of statistical analysis of single fibre proteomics is provided in detail in methods section. n = 5 participants per group.

Interestingly, in type I muscle fibres, we identified 136 differentially expressed proteins between controls and T2DM patients. Importantly, a large amount of sarcomeric proteins such as MYLK2, MYBPC2 and MYL4 were downregulated in T2DM muscle fibres (Fig. 4c, Table S2). On the other hand, in type II myofibres, controls and T2DM patients exhibited we observed 32 differentially expressed proteins between controls and T2DM group. Among these, mitochondrial-associated proteins, such as NDUFS5, NDUFA10 and NDUFS8 were downregulated in T2DM (Fig. 4d, Table S2). In summary, our analysis revealed distinct patterns of protein expression changes in type I and type II muscle fibres between control and T2D patients.

## Discussion

In the present study, we aimed to identify whether resting myosin biochemical and structural states were altered in T2DM. We observed a higher proportion of myosin heads in SRX and OFF state in T2DM patients than in controls. Interestingly, these alterations were only present in type I, slow oxidative, muscle fibres. The exact mechanisms underlying the increase in myosin super-relaxation may be complex but are likely to involve T2DM-specific hyper-glycated residues on the coiled coil region of myosin heavy chain type I (*MYH7*). Indeed, when acute hyper-glycation was induced in isolated single muscle fibres, the percentage of myosin heads in SRX increased. Besides hyper-glycation, myosin super-relaxation in T2DM may also involve a remodelling of sarcomeric proteins as revealed by our single-fibre proteomics analyses.

The discovery of SRX has remarkably shifted the field of muscle cell biology and has implied a potential direct link between resting myosin conformation and the metabolic rate of skeletal muscle (3, 4). However, much of the early work on resting myosin head biophysics has been on cardiac diseases, in particular, inherited hypertrophic cardiomyopathy (13, 14, 24). It is only recently that we have demonstrated that myosin biochemical states are dynamic and greatly influenced by non-genetic factors (25, 26). Based on these findings, investigating the potential involvement of changes to the dynamics of myosin in cardiometabolic syndromes such as T2DM was an obvious next step. Our primary finding, a higher fraction of myosin molecules in SRX and OFF state in T2DM, is striking. Critically, this means that the type I fibres of patients with T2DM have a reduced ATP turnover and thus energy consumption per muscle fibre. As skeletal muscle makes up around 40% of whole-body mass, even small changes in the energy consumption of this tissue can have significant consequences for whole-body energy expenditure (27). This is particularly interesting as patients with T2DM have been demonstrated to lose weight to a lesser extent than overweight, non-diabetic, controls, even when placed on matched weight management interventions (28). Future studies in this field should focus on the whether resting myosin SRX/OFF state is altered in obesity without the background of T2DM and whether dynamic changes to myosin conformation occur following weight loss and/or gain. Considering these findings, it is tempting to suggest that restoring SRX in the context of metabolic syndromes would promote ATP consumption of muscle and may favour a greater whole-body energy expenditure. This would then position myosin as a promising target for treating T2DM. Pharmacological compounds which can directly increase the DRX, specifically in skeletal muscle, may therefore hold therapeutic potential. It is important for future studies to investigate if differences in the myosin conformation exist between sexes as in the present study on male subjects were included in the T2DM patient cohort. This is of particular importance when considering that it has been shown that estradiol is able to disrupt the SRX (29).

Interestingly, the higher myosin super-relaxation only occurred in type I muscle fibres. Type I and type II fibres differ metabolically, with type I having a higher density of mitochondria and thus favoring oxidative respiration, whilst type II fibres have a relatively higher glycolytic capacity (30). Importantly, type I muscle fibres have a higher expression of the GLUT4 glucose transporter and have a far higher glucose handling capacity compared to type II fibres (31). Considering that glucose handling of skeletal muscle is one of the critical processes to be dysregulated during the development of T2DM, we believe that delineating whether myosin conformation and glucose handling in T2DM are biologically linked would be essential to pursue in future studies.

The molecular mechanisms underlying the changes in resting myosin conformations may be complex, but our study highlights the potential role of post-translational modifications such as glycation. Previous studies from Papadaki *et al*., showed that in T2DM patients, proteins of the cardiac myofilament were hyper-glycated, and that could reduce cardiac contractility (21, 32). In the SRX/OFF state, the myosin head is folded backwards and thus able to form interactions with this particular region (8, 24). The functional and experimental relevance of these unusual post-translational modifications was demonstrated using MG (33). When skeletal muscle fibres were treated with MG, we only observed an increased amount of myosin heads in the SRX state for type I muscle fibres of controls (not for T2DM). It is then highly plausible that, as a non-reversible covalent attachment of a carbohydrate or an *α*-oxo-aldehyde to an amino acid residue, the MG-treated myofibres from T2DM patients were already glycated/saturated prior to the incubation and thus were not significantly affected.

The proteomics analysis displayed differences in the expression of major sarcomeric proteins in type I skeletal muscle fibres of T2DM. Downregulated proteins included MYLK2 and MYL4. MYLK2 is a kinase which in adults is expressed specifically in skeletal muscle, although inherited pathogenic mutations to its associated gene lead to cardiomyopathies (34). Interestingly, a decreased expression in MYLK2 has previously been reported in individuals with high levels of insulin sensitivity (35). Future investigation into whether MYLK2 plays a functional role in the regulation of resting myosin conformation is essential, particularly due to the high level of regulation of myosin by PTMs. As a skeletal muscle specific protein, MYLK2 could be a promising target for the modulation of resting myosin states in T2DM and metabolic disease in a way which would not impact the adult heart.

In our analysis of type II fibres, we did not observe changes to sarcomeric proteins. Instead, type II fibres demonstrated differential expression of proteins associated with the mitochondria. In particular, we observed significant changes to the expression of NADH:ubiquitone oxioreductase subunit proteins, including NDUFS8, NDUFS5 and NDUFA10. These proteins have been previously demonstrated to be differentially expressed in the skeletal muscle of patients with T2DM (36). As this is the first time in which single fibre proteomics has been utilized in the investigation of T2DM this is the first identification of these proteins having specifically altered expression in type II fibres. Future experiments with fibre type specific investigation of oxidative phosphorylation in individuals with and without T2DM would be of benefit to the field as it would help define whether mitochondrial dysfunction is fibre-type specific in T2DM.

## Conclusion

Our findings indicate that T2DM patients undergo a remodelling of their resting myosin SRX/OFF state in type I muscle fibres, leading to a lower ATP demand. Such (mal)adaptation may be related to aberrant glycation levels on myosin heavy chains and may also be related to changes in the expression of sarcomeric proteins. These changes offer promising avenues for drug discovery as they suggest that if myosin biochemical and structural states were to be targeted pharmacologically and restored in T2DM, they would potentially help increasing the overall muscle/whole-body ATP consumption and thereby promote metabolic health.

## Methods

All these experimental approaches and statistics are presented in the Supplementary file.

## Data availability

The raw mass spectrometry data are deposited to the ProteomeXchange Consortium via the PRIDE partner repository with the dataset identifier PXD053022 (37).

## Acknowledgements

This work was generously funded by the Lundbeckfonden (R434-2023-311) to J.O. The X-ray experiments were under approval of the SPring-8 Proposal Review Committee (2022A1069). Mass spectrometry-based proteomics analyses were performed by the Proteomics Research Infrastructure (PRI) at the University of Copenhagen (UCPH), supported by the Novo Nordisk Foundation (NNF) (grant agreement number NNF19SA0059305). The CNIC is supported by the MCIN, the Instituto de Salud Carlos III, and the Pro-CNIC Foundation and is a Severo Ochoa Center of Excellence (grant number CEX2020-001041-S funded by MCIN/AEI/10.13039/501100011033).

## Data Availability

The data that support the findings of this study are available from the corresponding author upon reasonable request.

