## Supplementary Information for "Dysregulated Skeletal Muscle Myosin Super-relaxation and Energetics in Type II Diabetes"

**This PDF file includes:**

Methods
Tables S1 to S2
Figure S1
SI References

### Methods

#### Sample Collection

Human *vastus lateralis* biopsies were collected from two different studies. The collection of muscle biopsies from T2DM participants in Denmark was approved by the ethical committee for the Capital Region of Denmark (H-15010122) and was conducted in accordance with the declaration of Helsinki. All study participants were informed and signed a consent from before enrollment in the study. Characteristics from this study have been previously published and the biopsy procedure is described (1). For the collection of T2DM muscle biopsies in Canada, all procedures were approved by the Ethics Committee of the Université du Québec à Montréal (CIEREH-2020-3477). Informed consent was obtained from all participants. Skeletal muscle biopsy samples were obtained from the vastus lateralis muscle using a suction modified Bergström needle performed under local anesthesia. Samples were immediately frozen in liquid nitrogen and stored at -80C until use.

#### Single muscle fibre preparation

Cryopreserved muscle samples were dissected into small sections and immersed in a membrane-permeabilizing solution, described previously, (relaxing solution containing glycerol; 50:50 v/v) for 24 hours at -20°C, after which they were transferred to 4°C (2). Additionally, the rigor buffer for Mant-ATP chase experiments contained 120 mM potassium acetate, 5 mM magnesium acetate, 2.5 mM $K_2HPO_4$, 50 mM MOPS, 2 mM DTT at pH of 6.8. These bundles were kept in the membrane-permeabilizing solution at 4°C for another 24 hours. After these steps, bundles were stored in the same buffer at -20°C for use within two weeks.

#### Mant-ATP chase assay

Permeabilized skeletal muscle bundles were transferred to the relaxing solution and individual muscle fibres were isolated. Each muscle fibre was first incubated for five minutes with a rigor buffer. A solution containing the rigor buffer with added 250 µM Mant-ATP was then flushed and kept in the chamber for five minutes. At the end of this step, another solution made of the rigor buffer with 4 mM ATP was added with simultaneous acquisition of the Mant-ATP chase.

For fluorescence acquisition, a Zeiss Axio Scope A1 microscope was used with a Plan-Apochromat 20x/0.8 objective and a Zeiss AxioCam ICm 1 camera. Frames were acquired every five seconds with a 20 ms acquisition/exposure time using at 385 nm, for five minutes. These data were then fit to an unconstrained double exponential decay using Graphpad Prism 9.0:

$$\text{Normalized Fluorescence} = 1 - P1\,(1-\exp(-t/T1)) - P2\,(1-\exp(-t/T2))$$

Where P1 (DRX) is the amplitude of the initial rapid decay approximating the disordered-relaxed state with T1 as the time constant for this decay. P2 (SRX) is the slower second decay approximating the proportion of myosin heads in the super-relaxed state with its associated time constant T2.
Mant-ATP Chase Experiments were performed at ambient lab temperature (20°C) for all samples. Each isolated myofibre was then stained with a *MYH7* antibody (specific to type I MyHC) to decipher if the fibre was either a type I or type II fibre. Comparisons between muscle fibres sampled from different patient groups were then separated accordingly.

#### Synthesis and purification of methylglyoxal

MG was purchased from Sigma-Aldrich as a solution at 40 % in $H_2O$ (M0252). This solution was then further purified by steam distillation. The collected fractions were adjusted to pH 7.0 with NaOH and aliquots of them were reacted with $H_2O_2$ in order to indirectly quantify the pure MG concentration by titration with $KMnO_4$ (3).

#### Acute glycation Mant-ATP chase assay

Fibres were incubated for five minutes in a modified relaxing buffer solution containing 5.89 mM $Na_2ATP$, 6.48 mM $MgCl_2$, 40.76 mM propionic acid, 100 mM N,N-Bis(2-hydroxyethyl)-2-aminoethanesulfonic acid sodium salt, 6.97 mM EGTA, 14.5 mM sodium creatine phosphate dibasic and KOH to adjust the pH to 7.1 (4). After this incubation, fibres then went through the same protocol as described above of incubation in rigor buffer, Mant-ATP buffer and then fluorescence image acquisition. Following this, the same fibres were incubated in a modified relaxing buffer which contained 50 mM MG for 30 minutes to induce acute glycation of the proteins in these muscle fibres. This buffer contained 5.89 mM $Na_2ATP$, 6.48 mM $MgCl_2$, 40.76 mM propionic acid, 100 mM N,N-Bis(2-hydroxyethyl)-2-aminoethanesulfonic acid sodium salt, 6.97 mM EGTA, 43.5 mM sodium chloride 50 mM MG and KOH to adjust the pH to 7.4. Following this incubation these muscle fibres were incubated for five minutes in the modified relaxing buffer described above and then went through the exact same process of incubation of in rigor buffer, incubation with Mant-ATP buffer and then fluorescence image acquisition. For analysis purposes the results for each individual muscle fibre were matched before and after MG incubation to observe the effects of acute glycation upon myosin dynamics.

**X ray diffraction recordings and analysis**
Thin muscle bundles were mounted and transferred to a specimen chamber in relaxing buffer and then clamped at a sarcomere length of 2.00 µm. Subsequently, X-ray diffraction patterns were recorded at 15°C using a CMOS camera (Model C11440-22CU, Hamamatsu Photonics, Japan, 2048 x 2048 pixels) in combination with a 4-inch image intensifier (Model V7739PMOD, Hamamatsu Photonics, Japan). The X-ray wavelength was 0.10 nm, and the specimen-to-detector distance was 2.14 m. For each preparation, approximately 20-50 diffraction patterns were recorded at the BL40XU beamline of SPring-8 and were analysed as described previously (5). To minimize radiation damage, the exposure time was kept low (0.5 or 1 s) and the specimen chamber was moved by 100 µm after each exposure. Following X-ray recordings, background scattering was subtracted, and the major myosin meridional reflection intensities/spacing were determined as described elsewhere previously (5).

**Single Fibre Proteomics**
Proteomics measurements were conducted using a previously described workflow, in brief: an Evosep One HPLC system (Evosep) coupled via electrospray ionization to a timsTOF SCP mass spectrometer (Bruker) was utilized as the LC-MS system of choice (6). Peptides were separated using the '60 samples per day' chromatographic method before electrospray ionization by a Captive Spray ion source and a 10 µm emitter directly into the MS instrument. Samples were then measured using a DIA-PASEF workflow described elsewhere. Processing of the raw MS spectra was conducted using DIA-NN (version 1.8) in library-based mode, based on an in-house muscle fibre-specific MS library comprising 5000 proteins (7). The DIA-NN settings were as follows: double-pass mode was selected as the neural network mode, the "Robust LC (high accuracy)" was chosen as quantification strategy, prototypic peptides were selected for quantification together with enabling the match between runs option and setting the precursor FDR control to 1%. Unless specified, the other parameters remained as default settings.

**Single Fibre Proteomics data analysis/statistical testing**
Data analysis was performed using the R software (version 4.3.2) and multiple packages from the R environment. The protein groups matrix was loaded into the software and proteins were log2 transformed before being filtered for at least 70% valid values in at least one of the tested groups (T2DM or control). Next, the filtered protein matrix followed the limma (v 3.54.2) workflow, which included quantile normalization of all samples and differential expression testing between groups in a pseudobulk manner (8). To adjust p-values for multiple comparison testing, we applied the Benjamini-Hochberg correction for the *type I – type II* comparison, whereas the Xiao significance score was employed for the fibre type-specific comparisons of *controls – T2DM*, which takes into consideration both expression fold change and statistical significance. Therefore, proteins displaying a Xiao score under 0.05 were considered as differentially expressed between groups (9).

**Myosin heavy chain band Isolation for post-translational identifications**
Muscle biopsy samples were cut into 15 mg sections and immersed into a sample buffer (0.5 M Tris buffer at pH of 6.8, 0.5mg/ml bromophenol blue, 10% SDS, 10% glycerol, 1.25% mercaptoethanol) at 4°C. Samples were then homogenized and centrifuged allowing the supernatant to be extracted and used for SDS-PAGE gels (with stacking gel made with acrylamide/bis 37.5:1 and separation gel made with acrylamide/bis 100:1). Proteins were separated and individual MYH7 band was excised manually (10).

**Mass spectrometry-based glycation peptide mapping**
SDS-PAGE bands corresponding *MYH7* from control or diabetic groups were subjected to in-gel-digestion. After reduction with DTT (10mM) and alkylation of Cys groups with iodoacetamide (50mM), modified porcine trypsin (Promega) was added at a final ratio of 1:20 (trypsin-protein). Digestion proceeded overnight at 37ºC in 100mM ammonium bicarbonate at pH 7.8. Resulting tryptic peptides were loaded and washed on Evotips and separated in an Endurance 15 cm x 150 µm ID, 1.9 µm beads-EV1106 column using an Evosep one

HPLC system coupled to an Orbitrap Eclipse Tribrid mass spectrometer (Thermo Fisher, San José, CA, USA) and a 30 SPD preprogramed gradient.

MS analysis was performed using the data-independent scanning (DiS) method as described (11), with some modifications. Each sample was analyzed in a single chromatographic run covering a mass range from 390 to 1000 m/z. The DiS cycle consisted of 255 sequential HCD MS/MS fragmentation events with 2.5 m/z-windows from 390 to 900 m/z, and with 4 m/z-windows from 900 to 1000 m/z. HCD fragmentation was performed using a 33 normalized collision energy. MS/MS scans were performed using 70ms injection time, $3x10^5$ ions AGC target setting and 17,500 resolutions. The whole cycle lasted a maximum of 18 seconds depending on ion intensity during chromatography. The narrow windows used for fragmentation allowed peptide identification using conventional DDA searching algorithms. SEQUEST HT was used as implemented in Proteome Discoverer 2.5 (Thermo Scientific) against a database containing human myosins and with the following parameters: 2 maximum missed trypsin cleavage sites, 3 Da precursor mass tolerance and 30 ppm fragment mass tolerance. Oxidation in Met (+15.995 Da), CEL (+72.021 Da) and Carboxymethyl (+58.005 Da) in Lys, and MG-Hs (+54.011 Da) in Arg were set as variable modifications. Carbamidomethylation (+57.021 Da) in Cys was set as fixed modification (12).

**Statistical analysis**
Data are presented as mean ± standard deviation. Graphs were prepared and analyzed in Graphpad Prism v9. Relevant non-parametric statistical testing was utilized where data was unevenly distributed. Statistical significance was set to $p < 0.05$ unless otherwise stated.

**Table S1. Patient and control muscle biopsy samples used.**
Mean subject characteristics for the participants in the study of which biopsies samples were obtained from. Shown parameters are age (years), BMI (kg.m$^{-2}$) and myosin heavy chain (MyHC) breakdown (%) for all participants. Data is displayed as mean ± SD. Bilateral Student's t-test was used to calculate statistical significance. * = $p<0.05$ between groups. n = 9-11 subjects per group.

| | Control | T2DM |
|---|---|---|
| **Age (Years)** | 67.9 ± 12.9 | 67.9 ± 11.4 |
| **BMI (kg.m$^{-2}$)** | 29.76 ± 2.7 | 29.91 ± 2.4 |
| **MyHC Type I (%)** | 50.21 ± 9.46 | 43.93 ± 12.55 |
| **MyHC Type II (%)** | 49.79 ± 9.46 | 53.89 ± 12.55* |

**Table S2. Quantification of Identified Proteins from Single Fibre Proteomics.**
Detected protein abundance is listed for each individual muscle fibre analyzed. Significance levels are reported for each differential analysis performed. These are presented in the enclosed excel spreadsheet.

**Figure S1. Vseq characterization of AGE-modified peptide isoforms detected in MYH7.**
For each indicated peptide sequence the Vseq characterization is demonstrated.

**a**

AK$^{CEL}$NALAHALQSAR

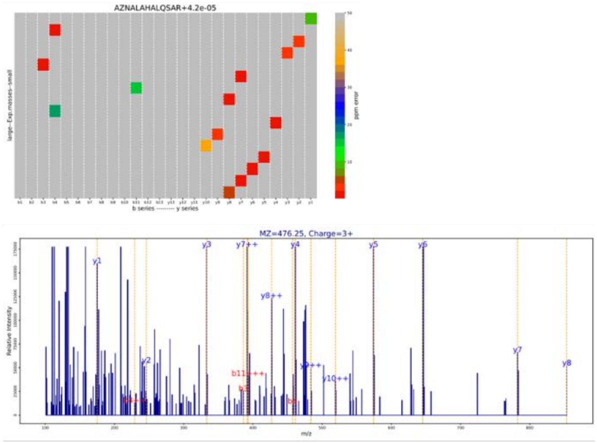


**b**

K$^{CEL}$MEGDLNEMEIQLSHANR

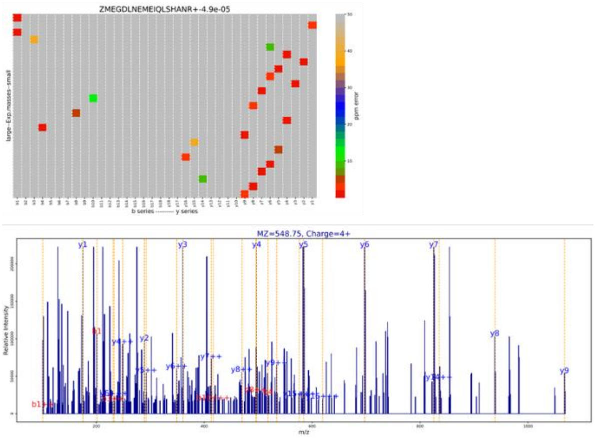


**c**

R$^{MGH1}$IK$^{Carboxymethyl}$ELTYQTEEDRK

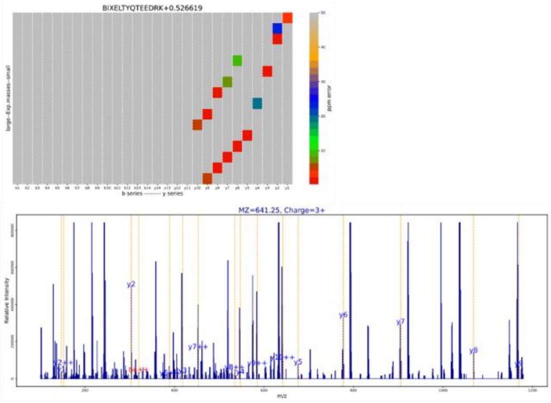

**d**

LAQR$^{\mathrm{MGH1}}$LQEAEEAVEAVNAK

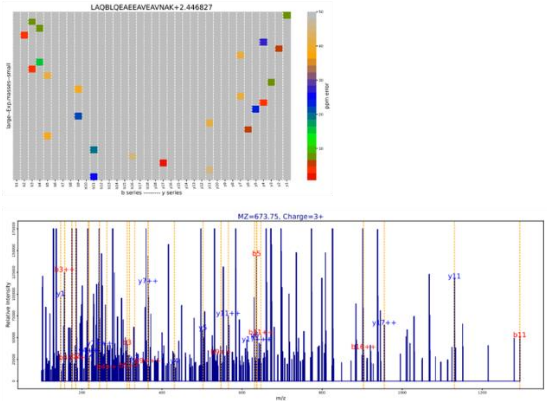


**e**

NHLR$^{\mathrm{MGH1}}$VVDSLQTSLDAETR

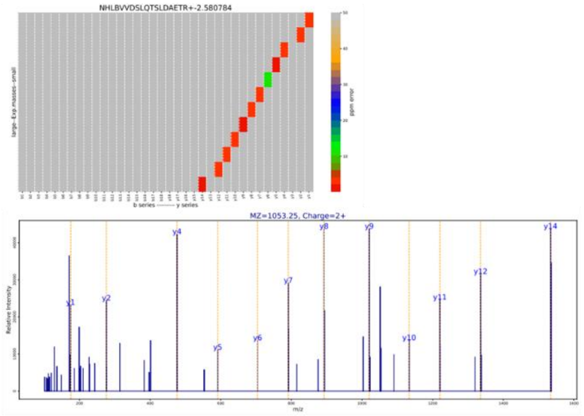


**f**

SR$^{\mathrm{MGH1}}$K$^{\mathrm{CEL}}$LAEQELIETSER

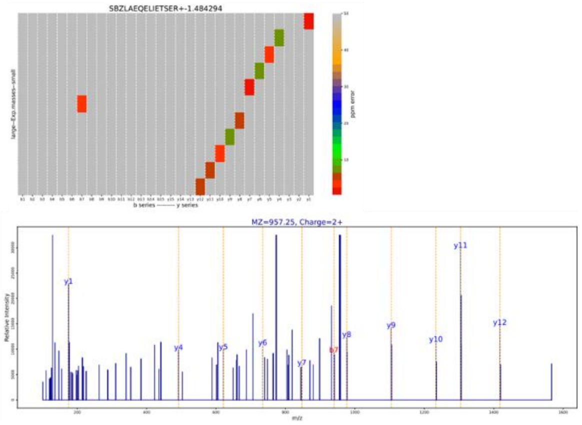

**g**

YR$^{MGH1}$ILNPAAIPEGQFIDSR

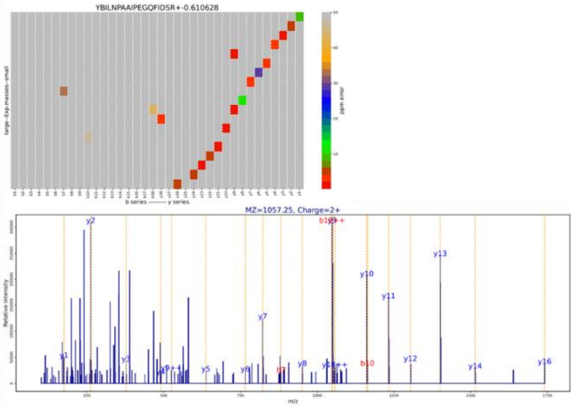
